# Adar-mediated A-to-I editing is required for establishment of embryonic body axes in zebrafish

**DOI:** 10.1101/2021.08.26.457081

**Authors:** Katarzyna Niescierowicz, Leszek Pryszcz, Cristina Navarrete, Eugeniusz Tralle, Marta Elżbieta Kasprzyk, Karim Abu Nahia, Katarzyna Misztal, Matthias Bochtler, Cecilia Winata

## Abstract

Adenosine deaminases (ADARs) catalyze the deamination of adenosine to inosine, also known as A-to-I editing, in RNA. Although A-to-I editing occurs widely across animals, and is well studied, new biological roles are still being discovered. Here, we study the role of A-to-I editing in early zebrafish development. We demonstrate that Adar, the zebrafish orthologue of mammalian ADAR1, is essential for establishing the antero-posterior and dorso-ventral axes and patterning. Genome-wide editing discovery revealed pervasive editing in maternal and the earliest zygotic transcripts, the majority of which occurred in the 3’-UTR. Interestingly, transcripts implicated in gastrulation as well as dorso-ventral and antero-posterior patterning were found to contain multiple editing sites. Adar knockdown or overexpression affected gene expression and global editing patterns at 12 hpf, but not earlier. Our study established that RNA editing by Adar is necessary for the earliest steps of embryonic patterning along the zebrafish antero-posterior and dorso-ventral axes.

## Introduction

RNA editing is a phenomenon of post-transcriptional alteration of transcript primary sequence ^[1]^. Its most common form is the A-to-I editing, in which adenosine (A) at the C6 position is deaminated, giving rise to an inosine (I) ^[2,3]^. As inosine pairs like guanine with tRNAs, A-to-I editing has the potential to alter the coding capacity of mRNAs, in some cases with drastic biological consequences ^[4,5]^. However, because of a prevalence of A-to-I editing in double-stranded RNA regions, most editing events do not affect the coding capacity of the genome.

A-to-I editing occurs widely in animals, from the earliest-diverging eumetazoan phyla to man ^[6,7]^. In vertebrates ^[8]^ and invertebrates ^[9]^, A-to-I editing prevents autoimmunity that is triggered by endogenous dsRNA ^[10–12]^. Another recurring theme of A-to-I editing is its role in the brain. In the mouse and zebrafish, editing of the GluR2 transcript is important for normal development of the nervous system ^[5,13,14]^. In the fruit fly, perturbed A-to-I editing causes behavioral phenotypes ^[15]^. In certain ant species, it determines caste-specific behavior ^[16]^. In the squid nervous system, extensive A-to-I editing is more prevalent in the giant axon system compared to the cell body, indicating region-specific editing within a neuron cell ^[17,18]^. In human and mouse, A-to-I editing contributes to germline integrity, by preventing the spread of Alu ^[19–21]^ and SINE elements ^[22]^, respectively. Higher editing prevalence in zebrafish testis and ovary compared to other organs ^[23]^ may hint to a role of A-to-I editing for germline integrity in non-mammalian vertebrates. However, to our knowledge, there is no evidence for this role in invertebrates yet.

A-to-I editing is catalyzed by adenosine deaminases (ADARs) ^[3]^. Most vertebrates have three paralogues that have arisen prior to vertebrate radiation, and can therefore be expected to have biochemically similar functions and substrate preferences ^[3,24]^. Apart from a C-terminal deaminase domain, all ADARs have at least two, and in some cases three double-stranded DNA (dsDNA) binding domains. ADAR1 additionally has a Z-DNA binding domain (ZBD) at the amino-terminal end of the protein that is missing from the other deaminases ^[25]^. Among the ADAR paralogues, only ADAR1 and ADAR2 are active, whereas ADAR3 has an inactive catalytic domain and appears to fulfill its biological role in the absence of catalytic activity ^[26,27]^.

In mammals, ADAR1 and ADAR2 are widely expressed, whereas ADAR3 is only expressed at low levels in the brain ^[3]^, particularly in the amygdala and hypothalamus ^[28]^. The majority of A-to-I editing is performed by ADAR1 and prevents dsRNA mediated autoimmunity ^[11]^. ADAR1/Adar1 occurs as two isoforms, known as p110 and p150 ^[29]^. Both share the deaminase domains, the dsRNA-binding domains, and one Z DNA-binding domain. Additionally, p150 possesses a second Z DNA-binding domain at the N-terminal end ^[30]^. Mice with a homozygous knockout of either both isoforms or only the p150 isoform cannot complete embryonic development. They die between embryonic day 11.5 and 12.5 from failed erythropoiesis and fetal liver disintegration ^[31,32]^, presumably due to stress in these cells ^[33]^. In contrast to *Adar1* null mice, *Adar2* mutant mice can complete embryonic development, but die subsequently from seizures, within three weeks of birth ^[5]^. Remarkably, this phenotype depends on a single editing event in the GluA2 AMPA receptor transcript. Even though multiple transcripts in the brain are edited ^[4]^, a point mutation in the GluA2 transcript suffices to suppress the Adar2 null phenotype. In humans, ADAR2 is associated with epilepsy, neurodegeneration, and autism ^[34]^.

In zebrafish, almost all data about A-to-I editing are descriptive. In contrast to most other vertebrates, which have three ADAR paralogues, zebrafish have four, due to a duplication of the ADAR2 ortholog into *adarb1a* and *adarb1b*. In the following, we use the official zebrafish nomenclature: *adar* for adar1, *adarb1a* for *adar2a, adarb1b* for *adar2b*, and *adarb2* for *adar3*. Transcripts for *adar*, and to a lesser extent *adarb1b*, are highly abundant during the first few hours of development. Transcripts of the other adenosine deaminase genes are scarce or absent during the first few hours. Although they are eventually expressed later on, the transcript levels remain lower than those of *adar* and *adarb1b* throughout development. Sequencing data suggests that editing in transcripts from repetitive genomic elements is pervasive in the first few hours of development, before the maternal to zygotic transition, and much less pronounced in later developmental stages ^[23]^. Editing in coding regions of genes sets in only later, roughly one day after fertilization. In adult organs, *adar* was most highly expressed in testis and heart, whereas *adarb1a* was highly expressed in heart and brain. Overall A-to-I editing was most pervasive in testes and ovaries ^[23]^. In contrast to the detailed information about the occurrence of editing, very little is known about functional consequences. It is known from earlier work that editing of GluR2 is conserved in zebrafish and is essential for normal development of the nervous system and cranial neural crest cells ^[14]^. However, it is unclear how conserved the role of A-to-I editing is otherwise, particularly for the early stages of development that differ greatly between mammals and zebrafish.

Here, we explore the role of A-to-I editing in early zebrafish embryos, focusing on the most highly expressed *adar*. Knockdown and overexpression experiments revealed that maternal *adar* is essential for zebrafish development, particularly during the earliest steps of antero-posterior and dorso-ventral patterning, and that this function is dependent on an intact deaminase domain. Transcriptome analysis uncovered prevalent A-to-I RNA editing during early embryogenesis, which affects transcripts known to play a role in gastrulation as well as dorso-ventral and antero-posterior patterning. *adar* knockout experiments further demonstrated later roles for the gene that could not be observed in the knockdown experiments because of the earlier lethality.

## Results

### The A-to-I RNA editing enzyme Adar and Adarb1b are expressed in the developing zebrafish embryo

To determine whether A-to-I editing activity exists during embryonic development, we revisited our transcriptome data ^[35]^ to check whether the enzymes responsible for A-to-I editing were expressed. In agreement with other recently published data ^[23]^, we detected transcripts of at least two deaminase paralogs, *adar* and *adarb1b*, from egg to 5.3 hpf (Fig. 1A). Transcripts of these two paralogs were present both maternally as well as zygotically, with *adar* being more abundant (more than 4-fold compared to *adarb1b* at each developmental stage). Moreover, we found that transcripts of both paralogs were consistently associated with polysome, starting from the egg stage up to 5.3 hpf (Fig. 1B). This finding suggests that these transcripts are constantly undergoing translation at developmental stages preceding and after the activation of zygotic genome ^[35]^. The observation that both paralogs were already expressed, and their transcripts associated with polysomes at egg stage suggests that RNA editing events occur prior to fertilization and may be crucial for early development. Interestingly, a substantial increase in *adar* expression occurs after the mid-blastula transition (MBT), suggesting that the role of this gene extends beyond the period of transcriptional silence in early embryogenesis (Fig. 1A). At larval stage, the expression of both gene paralogs was not spatially restricted although more abundant in the nervous system of the developing embryo as shown by whole mount *in situ* hybridization of *adar* (Fig.1C, D) and *adarb1b* (Fig. 1E, F) in 24 hpf zebrafish embryos.

**Figure 1.**
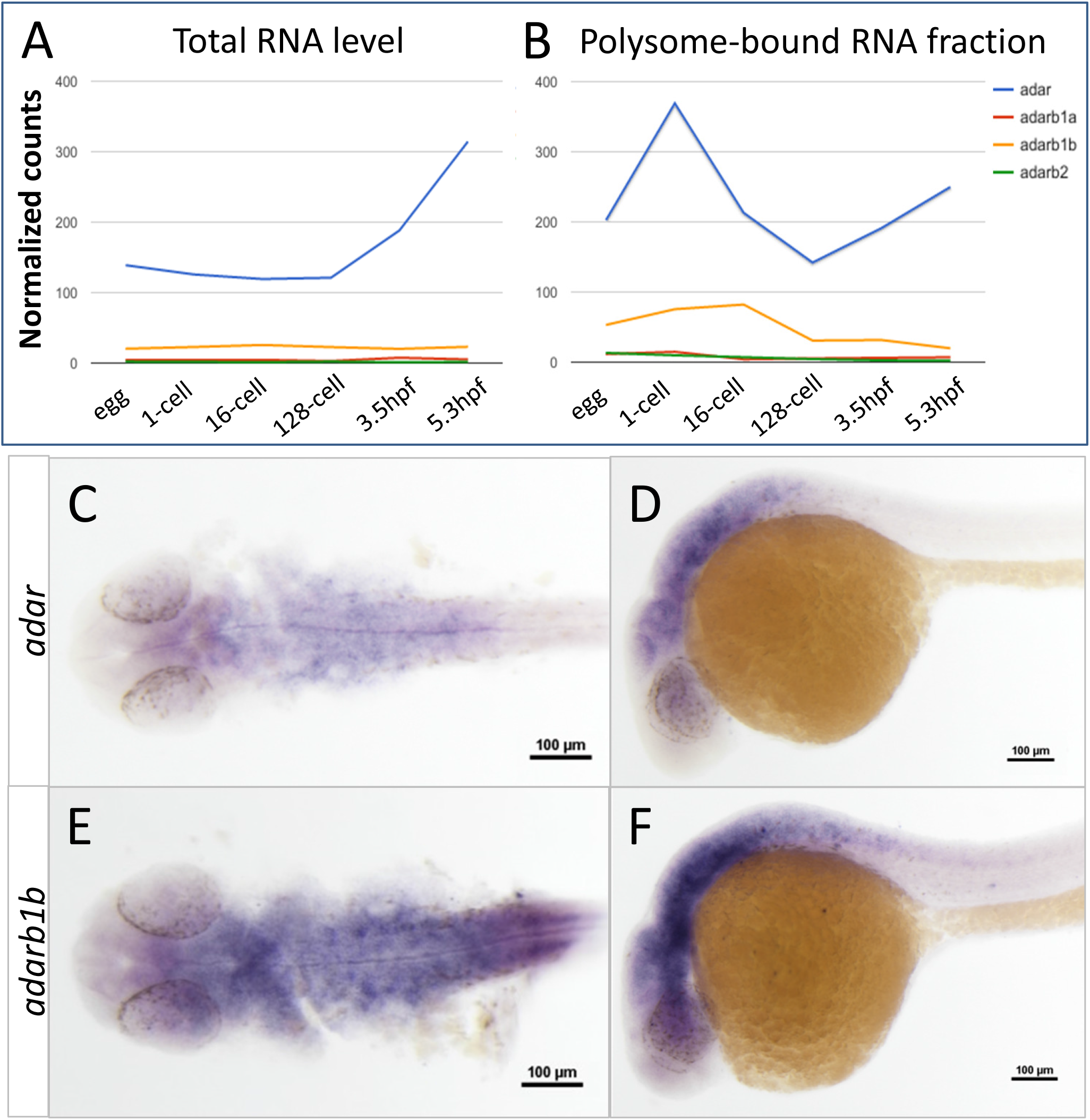
Spatiotemporal expression pattern of Adar family genes in zebrafish. (A) Expression of zebrafish *adar* family based on transcriptome profiling of developing embryos (RNA-seq) ^[54]^. Expression levels are plotted for developmental stages: egg, 1 cell, 16 cells, 128 cells, 3.5 hpf and 5.3 hpf. Levels of (A) total RNA and (B) polysome-associated RNA of four zebrafish *adar* paralogs are given. Whole mount *in situ* hybridization shows expression pattern of *adar* (C, D) and *adarb1b* (E, F) in 24 hpf zebrafish embryos.

### The A-to-I editing activity of Adar is required for early embryonic patterning

To investigate whether Adar plays a biological role in early zebrafish development, we used morpholino oligonucleotides (MO) to knock-down Adar, the highest expressed among all zebrafish ADAR paralogs (Fig. 1A). Adar MO-injected embryos developed a range of phenotypes which is initially evident at gastrulation and subsequently observed to affect posterior body axis by 24 hpf (Fig. 2A-C, H). The most severe phenotypes manifested in a lack of almost all posterior structures and crooked body axis (Fig. 2C). In addition, the notochord in morphant embryos was disorganized, with unevenly shaped and distributed vacuoles instead of the neat “stack of coins” arrangement in wild-type (Fig. 2B, C). This abnormal MO phenotype was dose-dependent and could be rescued with the wild-type *adar* mRNA, in which an increased percentage of embryos appeared normal or exhibited mild phenotype with proper body axis organization and tail length (Fig. 2D, H; Supplementary Fig. 1). On the other hand, knockdown with up to 2ng of MO against Adarb1b, which was expressed at a lower level in the early embryo, did not result in any observable phenotype (Supplementary Fig. 1). To verify whether the biological role of Adar depends on its RNA editing activity, we generated *adar* mRNA E1030A designed after a similar construct in mammals ^[36]^, in which the deaminase domain was mutated (Supplementary Figure 1). The E1030A mRNA was unable to rescue MO-induced phenotype in developing embryos, resulting in comparable number of abnormally developed embryos with severe posterior axis defects to that of MO-injected larvae (Fig. 2E, H). This suggests that a functional deaminase domain, catalyzing the A-to-I editing in dsRNA, is essential for early embryonic development. These results strongly suggest that the A-to-I editing activity of Adar is necessary for the specification of early embryonic axes.

**Figure 2.**
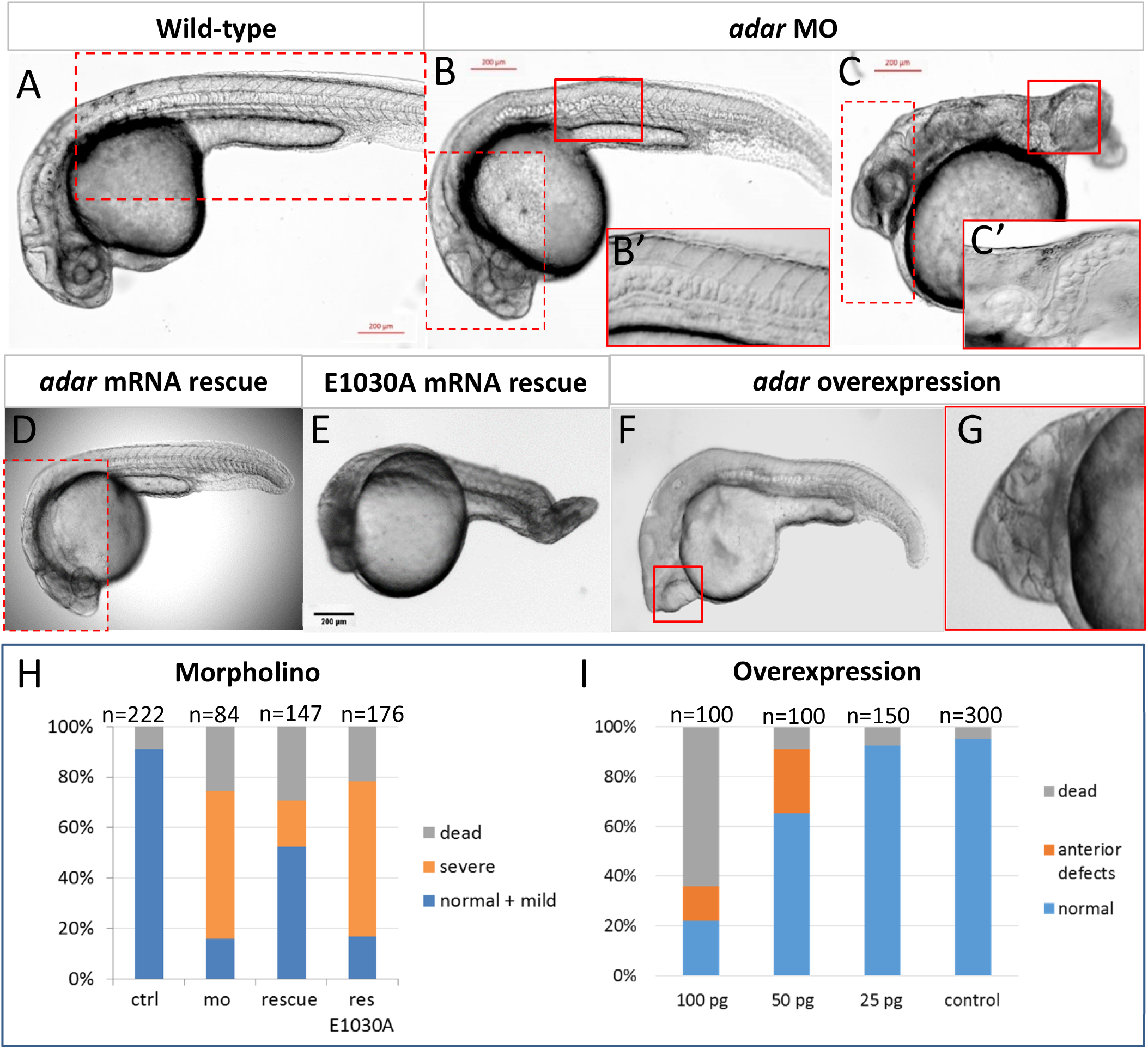
Phenotypic defects at 24 hpf caused by *adar* knockdown and overexpression. (A) Wild-type (B, C) Adar MO-injected embryos develop abnormal phenotype in the posterior part with disturbed body axis, shortened tail and crooked, disorganized notochord. (D) MO phenotype can be fully rescued with wild-type mRNA injection. (E) Mutant *adar* mRNA E1030A with inactivated editing domain could not rescue the malformed phenotype. (F,G) Phenotype defects caused by *adar* mRNA overexpression. The anterior defects, including cyclopia and head malformations are mRNA dose-dependent. Inset marked by red boxes denotes overlaid identical image taken at different focal plane. (H, I) Injection statistics of Adar MO, rescue, and mRNA overexpression.

Whereas Adar MO affected the development of the posterior part of the body, *adar* mRNA overexpression caused significant abnormalities of the anterior part, which include anomalous eye development, most often manifested in cyclopia, deformed cranium, and reduced or absent brain compartment, particularly at the anteriormost region (Fig. 2F, G, I). These defects were most prominently observed with 50 pg injection of mRNA. The effect of *adar* mRNA overexpression was dose-dependent with a 64% mortality for the 100 pg of injected mRNA, 9% for 50 pg and 7% for 25 pg (compared to wt which had 5% mortality). Collectively, our results show that Adar plays a key role in the earliest steps of embryonic patterning.

In order to characterize the observed morphological defects of Adar loss- and gain- of function in more detail, we assessed the expression of several marker genes indicating various embryonic structures. The most prominent phenotype of Adar disruption is that of the development of structures along the anteroposterior axis. In order to characterize the effect of Adar loss- or gain-of function on the development of anterior embryonic structures, we utilized the expression of *pax6* and *tbx2b* to collectively indicate, among which, the forebrain, midbrain, and hindbrain, anterior spinal cord, optic vesicle, and otic vesicle (Fig. 3A, D, G, J) ^[37] [38]^. In *adar* morphants where morphological defects were predominantly observed in the posterior region, these anterior structures were preserved and appeared similar to wild-type in terms of their size and organization (Fig. 3B, E, H, K). On the contrary, anteriormost brain regions and optic vesicle were indistinguishable in embryos overexpressing *adar* (Fig. 3C, F, I, L). In particular, *pax6* expression shows that the diencephalon and telencephalon expression domain were unrecognizable in *adar*-overexpressing embryos (Fig. 3C, F). Moreover, *tbx2b* expression domains of telencephalon, left and right optic vesicles, and the epiphysis appeared as a fused region, whereas the more posterior expression domains of the trigeminal ganglion and otic vesicles appeared less affected although hypomorphic compared to control (Fig. 3I, L). While the eye field is derived from the anterior neural plate and therefore arose as a consequence of antero-posterior patterning ^[39]^, its failure to split in the midline (cyclopia) is a hallmark of defect in the convergent-extension movement during gastrulation which is dependent on the establishment of the dorsoventral axis ^[40,41]^. The observed phenotype of *adar* overexpression therefore suggests that proper establishment of these two body axes was affected.

**Figure 3.**
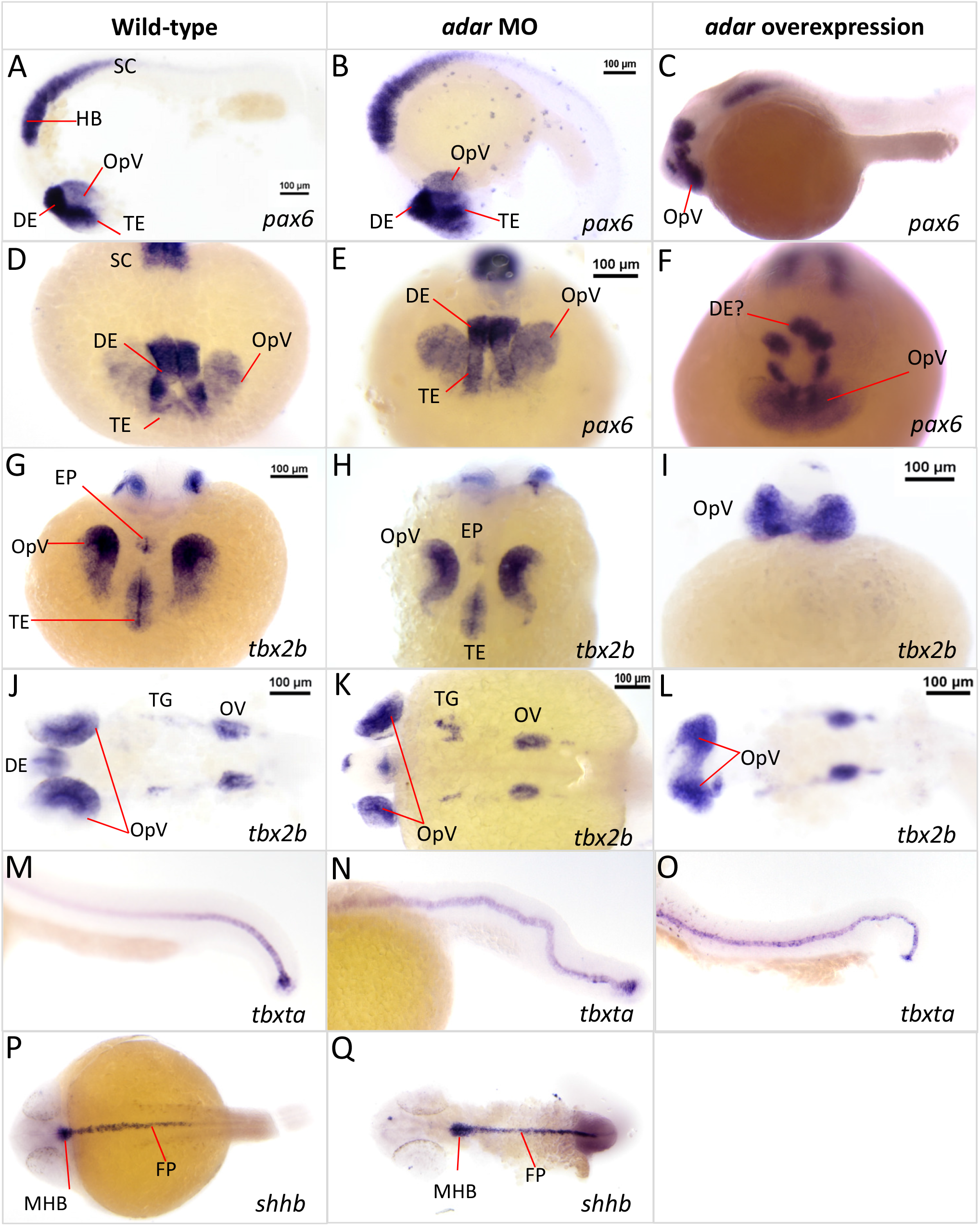
Embryonic patterning defects due to Adar loss- or gain-of-function. Whole mount *in situ* hybridization detection of marker genes in Adar morpholino knockdown and *adar* mRNA overexpression. (A-F) *pax6* expression demarcates anterior brain structures including optic vesicle (OpV), telencephalon (TE), and diencephalon (DE) as well as hindbrain and anterior spinal cord. (G-L) *tbx2b* expression marking TE, dorsal part of the OpV, DE, and otic vesicle (OV). (M-O) expression of *tbxta* marks the notochord. (P, Q) *shhb* expression indicates floor plate (FP) and midbrain-hindbrain boundary (MHB).

To examine the defects in posterior structures, we used as reporter genes *tbxta* and *shhb* which mark the notochord and floor plate neurons of the spinal cord, respectively (Fig. 3M, P). Similar to the anterior markers, *shha* and *tbxta* were expressed in the case of both *adar* loss- or gain-of-function, suggesting that the neural and midline cell identities were preserved (Fig. 3N, O, Q). However, the body axis including the notochord and spinal cord were crooked in *adar* morphants as well as in *adar* overexpressing embryos albeit to a lesser extent in the latter (Fig. 3N, O). Importantly, loss- or gain-of-function of *adar* do not appear to affect the specification of cell identity as evidenced by the preserved expression of *pax6, tbx2b, shha, and tbxta*. Rather, a more pronounced effect is seen on the gross anteroposterior and dorsoventral patterning of the embryo. Collectively, these observations suggest that Adar-mediated A-to-I editing may be involved in the patterning along the two early embryonic axes of the zebrafish.

### A-to-I RNA editing is prevalent in maternal and early zygotic transcripts during normal embryogenesis

The effects of Adar disruption on embryonic patterning led us to ask to what extent A-to-I editing occurs in early embryogenesis, and whether transcripts of genes involved in this process were specifically affected. To profile global A-to-I RNA editing in zebrafish we sequenced a trio of wild-type sample of both parental genomes and the transcriptome of their offspring at three developmental stages: 1.5 hpf (16-cells, pre-MBT), 3.5 hpf (high, MBT) and 5.3 hpf (50% epiboly, post-MBT) (Fig. 4A). Comparison of zygotic transcriptomes with the genomic sequence of parents allowed us to pinpoint RNA editing events by identifying mismatches between the genomic and transcriptomic reads. As Inosine (I) is structurally similar to a G due to the presence of the 6-oxo group, reverse transcriptase incorporates a C in the corresponding position during RNA-seq library synthesis; thus, in the original transcript strand, a G is inferred. Therefore an A-to-I editing event can be identified as an A-G mismatch between the parental genome and the corresponding embryonic transcript. Since the RNA-seq was unstranded, the same applies as well to a T-C mismatch. Strikingly, our analyses revealed a disproportionate enrichment of A-G and T-C mismatches compared to other possible base mismatches which could occur stochastically at all three developmental stages (Fig. 4B). This strongly suggests the presence of A-to-I RNA editing in maternally deposited as well as in zygotic transcripts. Altogether, we identified 44,007 RNA editing sites: 11,374 of which were shared by all three samples, 6352 between 1.5 hpf and 3.5 hpf, 1393 between 1.5 hpf and 5.3 hpf, and 1,564 between 3.5 hpf and 5.3 hpf. Apart from this, 6117, 5686 and 11,521 were specific to 1.5 hpf, 3.5 hpf and 5.3 hpf, respectively (Fig. 4C, Supplementary Table 1). These stage-specific editing events indicate differential patterns of RNA editing throughout early embryogenesis and suggest that this process may play a role in embryogenesis. Interestingly, merely 2% of RNA editing sites occurred within coding sequences (CDS), while the majority (38% in 1.5 hpf and 3.5 hpf and 27% in 5.3 hpf) occurred in 3’-UTR regions (Fig. 4D, Supplementary Table 1). A large fraction of editing was assigned as ‘genic_other’ due to overlap between intron/exon/UTRs from multiple transcripts. Finally, ~17% of editing events in each stage were detected in ‘intergenic sequence’, but this may be due to unannotated transcripts. The low abundance of A-to-I editing present in the coding regions (CDS), none of which resulting in missense mutations, ruled out the possibility that editing functions through expanding the coding repertoire of expressed transcripts by altering amino acid composition. We also observed plenty of editing occurring in DNA repeat regions with a notable increase in editing frequency in retrotransposons (LTR, LINE, and SINE) in 5.3 hpf compared to earlier developmental stages (Fig. 4E, Supplementary Table 2).

**Figure 4.**
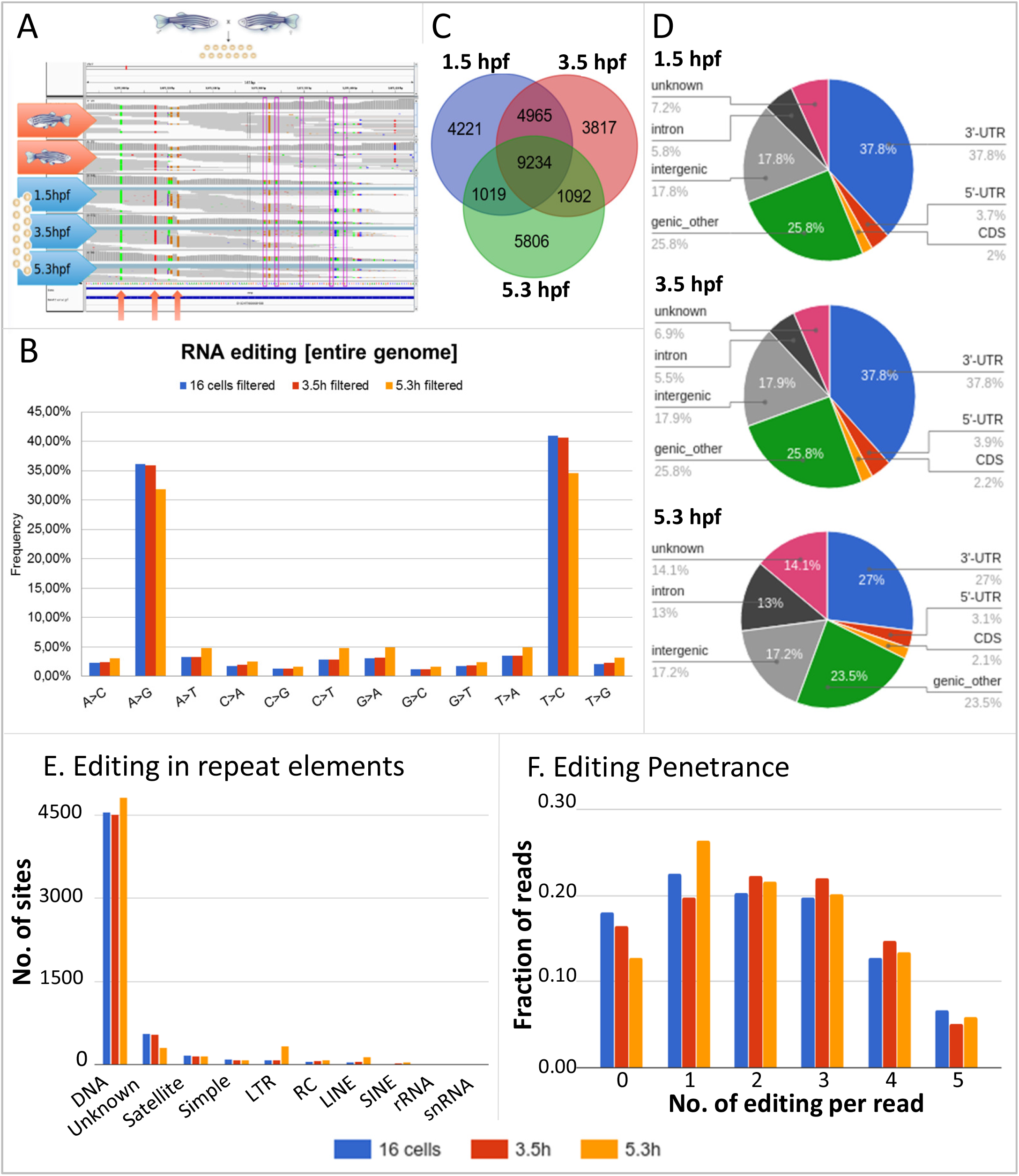
A-to-I RNA editing sites in early embryonic development. (A) Schematics of A-to-I RNA editing discovery through sequencing of a parent-offspring trio. The genome sequence of the parents are used as a reference set to distinguish between polymorphisms and editing. (B) Mismatches between RNA and DNA sequencing data. As RNA libraries were not strand selective, mismatches were read as their complement (i.e. T->C instead of A->G, or C->T as G->A) in roughly half of all cases. (C) Overlap of editing sites at different time points. The 1.5 and 3.5 hpf samples were more similar to each other than to the 5.3 hpf sample, probably because of replacement of maternal by zygotic transcripts at the MZT. (D) Association of editing sites with genomic features. A large fraction of RNA editing is classified as ‘genic_other’ due to overlap between introns/exons/UTRs from multiple transcripts. (E) Number of editing sites in transcripts stemming from different classes of repeat elements. (F) Number of editing events in individual reads encompassing 10 potential editing sites. The majority of individual reads contained 1-3 RNA editing sites, and never more than 5 editing sites.

To assess the penetrance of A-to-I editing on individual RNA molecules, we calculated the occurrence of editing events in RNA-seq reads. Based on alignment of 75bp reads, we observed a substantial number of reads with no editing at all for a given region (Fig. 4F). Interestingly, the number of RNA editing sites in a single read never exceeds 50% of the total number of editing events identified in the given transcript region (Fig. 4F, Supplementary Table 3). This suggests that A-to-I editing rarely occurs at all possible editing sites in a given RNA molecule, although sequencing with much longer reads is required to precisely determine the true penetrance. We then ask whether RNA editing events were associated with translation efficiency and/or RNA stability. Using our previously published polysome profiling dataset to determine translation rate ^[35]^, we found that transcripts undergoing more active translation tend to be less edited than those not associated with polysome (Supplementary Fig. 2). Interestingly, this is true only for editing events detected at post-MBT stages (3.5 hpf and 5.3 hpf), but not for pre-MBT (1.5 hpf). With regards to RNA stability, no obvious difference was observed in expression levels between low (1-9 sites) and high (more than 10 sites) edited transcripts when 1.5 hpf and 3.5 hpf stages were compared. However, highly edited transcripts undergo much lower expression change between 1.5 hpf and 5.3 hpf (Supplementary Fig. 3).

We sought to identify which genes were subject to A-to-I editing at each developmental stage. We identified 639, 634, and 562 genes having at least two editing sites at any position within their transcript at 1.5 hpf, 3.5 hpf, and 5.3 hpf respectively (Supplementary Table 4). Among these, 9, 9, and 10 genes had editing sites occurring in both 3’UTR and coding sequence at each respective stage. Interestingly, although no developmentally relevant GO terms were found to be significantly enriched among edited genes (Supplementary Table 5), we found several key genes known to be implicated in anteroposterior and dorsoventral patterning (Fig. 5). Of note, members of the Wnt signaling pathway were found to be edited. Several Frizzled receptors of Wnt signaling, *fzd3b, fzd5*, and *fzd8b* were edited at both maternal stages, while *fzd7b* were edited at 5.3 hpf. The Wnt downstream effector *tcf7l1b* were found to be consistently edited at all three stages. We also found two members of the FGF signaling pathway, *fgfr1a* and *extl3*, consistently edited at all three stages observed. Interestingly, *fgfr1a* is one of the highest edited transcripts, containing 99 edited sites in both its coding sequence and 3’-UTR. Mutation of FGFR1 in humans is associated with holoprosencephaly ^[42]^ which is reminiscent of the observed *adar* overexpression phenotype. Other genes which were consistently edited throughout the three stages include *furina* which plays a role in craniofacial development ^[43]^ and two genes involved in gastrulation, *dusp4* and *ezrb* ^[44,45]^. That transcripts of these genes were found to be edited throughout maternal and early zygotic stages suggests the role of A-to-I editing in regulating multiple aspects of anteroposterior and dorsoventral patterning.

**Figure 5.**
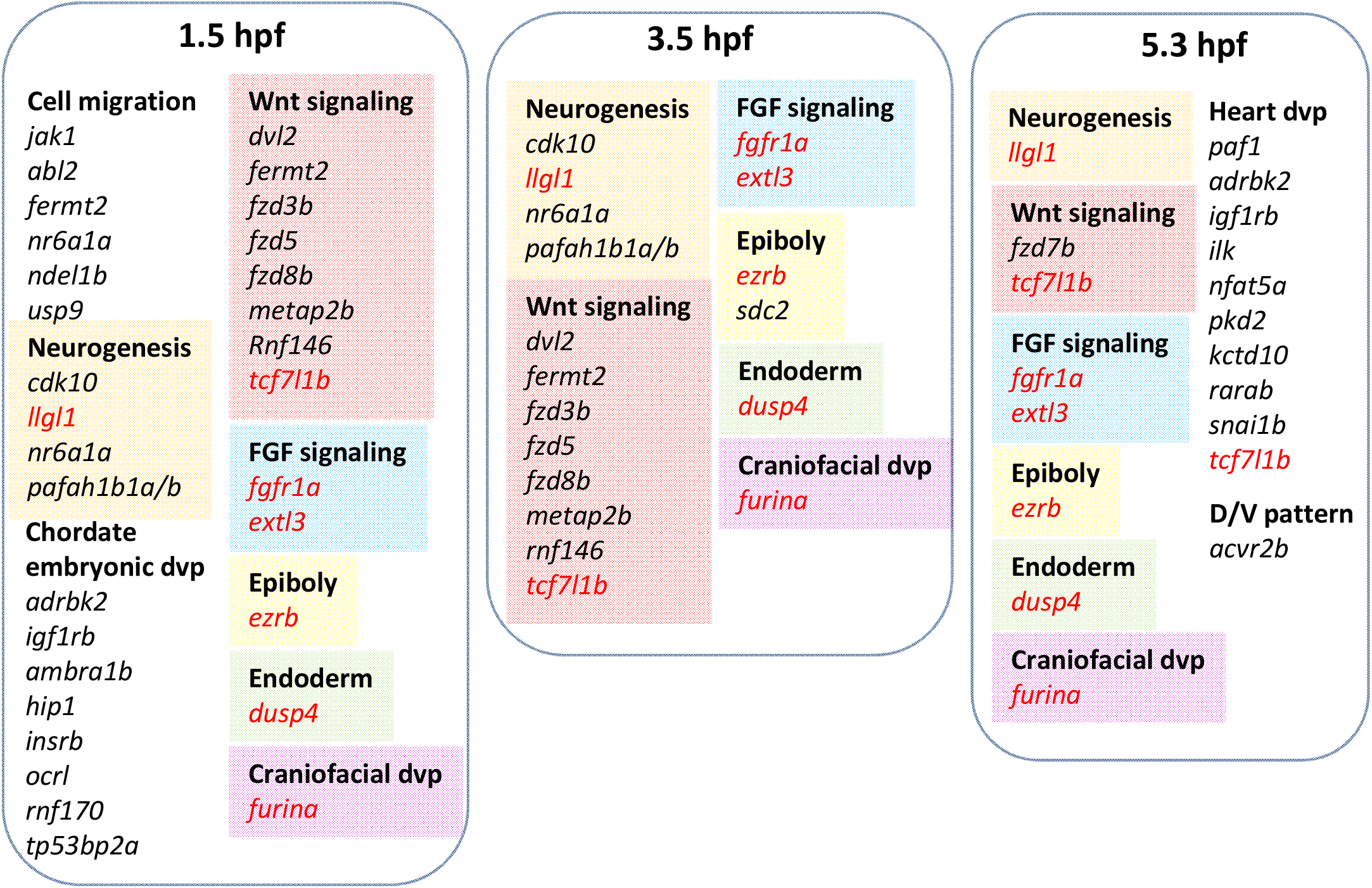
Selected transcripts of developmental signaling pathway genes implicated in dorsoventral and/or anteroposterior patterning containing two or more RNA editing sites detected at their 3’-UTR region. Gene names in red denotes those that are commonly edited at all three stages.

### Maternal-Zygotic Adar loss of function affects dorsoventral patterning without observable changes in global A-to-I editing patterns

To uncover the role of Adar at the molecular level, we performed transcriptome sequencing of wild-type, Adar MO knockdown (KD) and Adar overexpressing (OE) embryos at 128-cells (before ZGA), 5.3 hpf (after ZGA), and 12 hpf (when embryonic patterning is established). We hypothesized that alterations in the level of *adar* transcripts may evoke consequential differences in mRNA editing and/or changes in the expression of potential target genes. To this end, we performed comparative transcriptome analysis of wt vs OE and wt vs KD in each developmental stage. Surprisingly, no substantial changes in global A-to-I editing levels were found in KD and OE conditions at 128-cell and 5.3 hpf stages (Supplementary Fig. 4A). Moreover, during these two stages, samples clustered according to developmental stage (Supplementary Fig. 4B) and no substantial changes were observed in global gene expression profile (p<0.05; Supplementary Fig. 4C). Only 539 sites have a slightly elevated editing level in OE than in KD in both developmental stages (Supplementary Table 6).

In contrast, at 12 hpf, a more noticeable change in both global editing pattern and gene expression profile were observed between control and Adar gain- or loss-of-function at 12 hpf (Fig. 6A; Supplementary Fig. 5). Unlike at earlier stages, samples clustered according to conditions rather than developmental stage (Fig. 6B). Interestingly, in all three replicates, Adar KD caused a modest but more noticeable change in editing frequency compared to the earlier stages (Supplementary Fig. 5). Moreover, at 12 hpf, we observed 827 and 5054 genes differentially expressed in Adar KD and Adar OE respectively, compared to control (p<0.05; Supplementary Table 7). GO analysis (Supplementary Table 8) revealed that Adar KD generally caused the upregulation of genes regulating epiboly (*nanog, cacnb4b*), gastrulation (*bcl2l10, mylipa*), and ectoderm development (*pou5f3, cdh1*), while concurrently causing downregulation of those implicated in convergent-extension (*wnt11, creb1a, ppp1cb*) and mesoderm development (*tbx16, hes6, her7, mcdh2, myf5, msgn1*). On the other hand, Adar OE resulted in the upregulation of genes involved in the development of mesodermal structures (*myf6, bves, tcf21, tbx5a, apln, mef2aa*), while downregulating genes regulating epiboly (*chuk, epcam, mapkapk2a, cldne*), dorsoventral pattern formation (*sox11b, dusp6, ved, vox, bambia, acvr1ba, ctnnb2*), and brain development. Interestingly, out of the 383 genes common between the two conditions, 283 commonly downregulated genes in both Adar KD and OE included genes known for their role in anterior-posterior as well as dorsoventral patterning (Fig. 6C; Supplementary Table 7), such as *cdx4* ^[46]^, *szl* and *ved* required for DV patterning ^[47,48]^, and several *hox* genes ^[49]^. This suggests that both Adar KD and OE may act on factors with downstream consequence of suppression of these axis-regulating genes.

**Figure 6.**
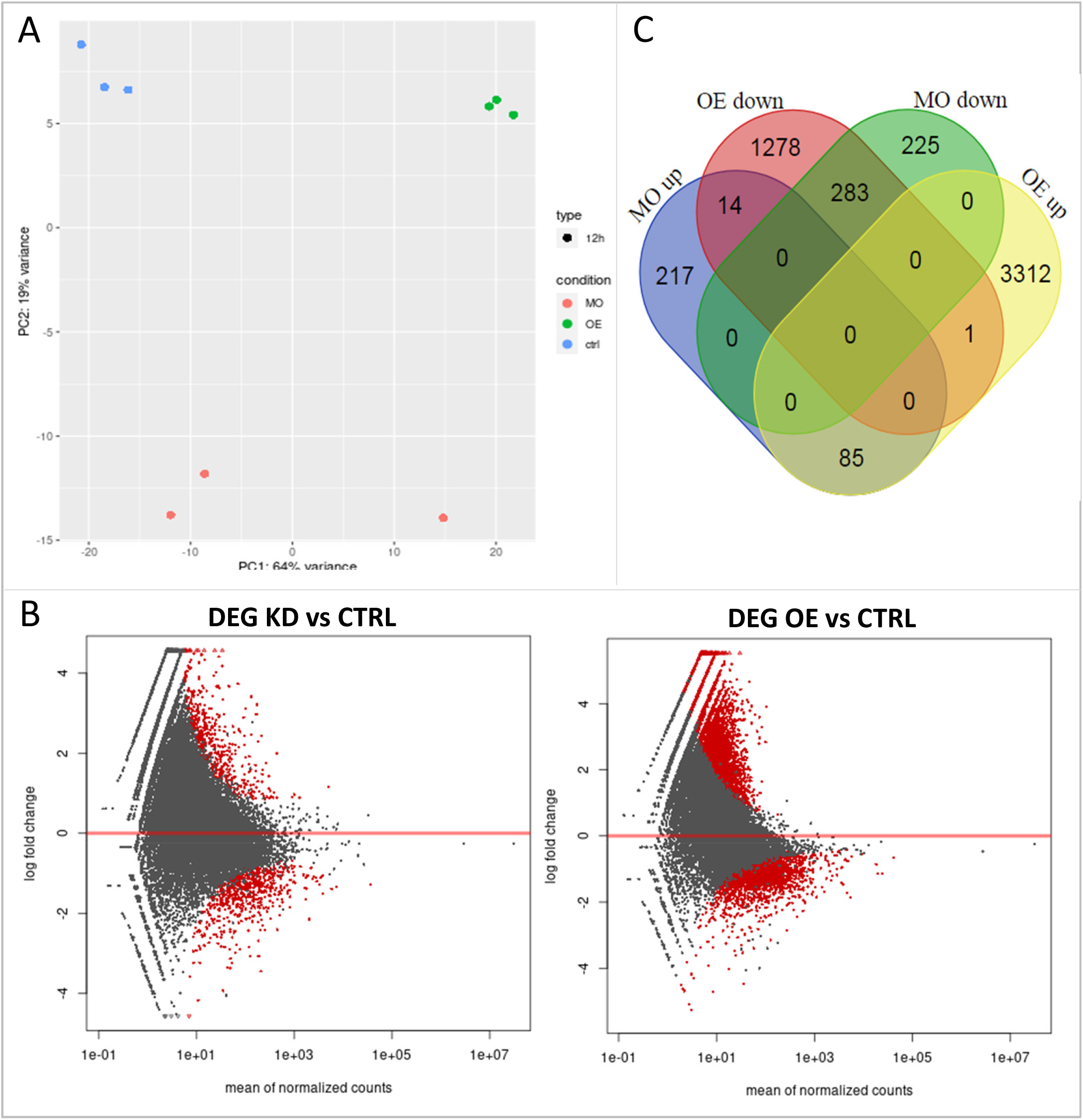
Differentially expressed genes at 12 hpf in Adar KD and OE: **(A)** Principal component analysis of control, Adar knockdown and overexpression samples based on their transcriptome profile. (B) Number of genes differentially expressed in each condition and their overlap. (C) Differential expression analysis of Adar KD and OP compared to control. Genes with significant change in expression (p<0.05) in red.

As Adar gain- and loss-of-function resulted in almost opposite defects in anterior-posterior and dorso-ventral patterning, we compared the transcriptome from the two conditions and found 1218 differentially expressed genes (Supplementary Table 7). Genes regulating the development of mesodermal structures (*bmpr2a, emilin3a, msgn1, gata1a, meis1b, etv2, pdgfra*) as well as *hox* family genes (*hoxa4a/9b, hoxb3a/5a, hoxc1a/3a/6a, hoxd4a*) were upregulated in OE compared to KD (Supplementary Table 8). On the contrary, genes regulating epiboly and gastrulation (*chuk, epcam, mapkapk2a, snai1a*), convergent-extension (*dsc2l, gpc4, tdgf1, ptpra, prmt1*), and dorsal-ventral patterning (*sox11b, ctnnb1, ppp4cb, tll1*) were downregulated in OE compared to KD (Supplementary Table 8). These observations agree with the phenotypes resulting from Adar disruption which affected structures along the anteroposterior and dorsoventral axes, where an excess of Adar affected dorsal and anterior structures while Adar deficiency affected more ventral and posterior structures. Taken together, the consequences of Adar disruption at the molecular level could be observed by 12 hpf, which affected the expression of genes involved in the development of various embryonic structures. This suggests that, although no significant changes were observable in global editing levels and transcriptome up to 5.3 hpf, Adar is necessary during the crucial period of embryonic patterning, between gastrulation and 12 hpf.

### Zygotic Adar is not essential for early embryogenesis

We then asked whether zygotic Adar activity is required for later events of embryogenesis. To this end, we created a zebrafish *adar* mutant line (*adar-/-*) using the CRISPR/Cas9-based gene knock-out method. A 5 bp deletion was introduced into the second exon of the *adar* gene, resulting in a premature stop codon and polypeptide consisting of 4% (39 aa) of the full length, functional Adar protein (917 aa) (Fig. 7A). To generate homozygous *adar* mutants we incrossed F1 heterozygous individuals and observed the F2 offspring. Surprisingly, no developmental phenotypes were observed in any of the offspring. We then raised these F2 individuals and genotyped them at approximately 3 months post-fertilization by fin clipping. Surprisingly, we found 76% heterozygous and 24% wild type fish among the 200 screened samples. None of the adult F2 individuals were homozygous for the *adar-/-* mutant allele (Fig. 7B). This suggests that the homozygous *adar-/-* knockout is not viable despite their lack of developmental abnormalities. To determine the fate of the homozygous *adar-/-* mutants, we genotyped F2 individuals (~300 individuals) at 3 dpf. Interestingly, the distribution of homo-, heterozygous and wild-type individuals was in agreement with the Mendelian ratio (Fig. 7B), indicating that mortality of the homozygous *adar-/-* individuals occurred later than 3 dpf and before adulthood. In order to pinpoint the timing of mortality, we followed the development of homozygous individuals until the point of death. The *adar-/-* larvae did not demonstrate any observable morphological abnormalities when compared to heterozygotes or wild-type individuals. However, significant mortality was observed between second- and third-weeks post fertilization. Nearly 100% of *adar*-/-mutants died within this period, with only one surviving until the third month post fertilization, with substantial growth impairment. To eliminate the possibility of genetic complementation ^[50]^ causing an attenuation of the *adar-/-* phenotype, we analyzed mRNA levels of *adar* as well as three other *adar* family members: *adarb1a, adarb1b* and *adarb2*, in wild-type and *adar-/-* homozygotes. No significant changes in the mRNA levels of *adar, adarb1a, adarb1b and adarb2* mRNAs were observed in *adar-/-* individuals when compared to wild-type at 3 dpf (Fig. 7C). Our findings therefore suggest that the lack of phenotypic alterations in *adar-/-* larvae, when compared to *adar* KD (MO-injected) was not due to genetic compensation. It is possible that the late mortality of zygotic *adar* KO stems from the presence of maternally deposited *adar* mRNAs and/or proteins from heterozygous mothers, which might be sufficient to drive early developmental processes in the first few days post fertilization. Altogether, these observations suggest that zygotic Adar is not required for early embryogenesis although its function is still essential for life later on.

**Figure 7.**
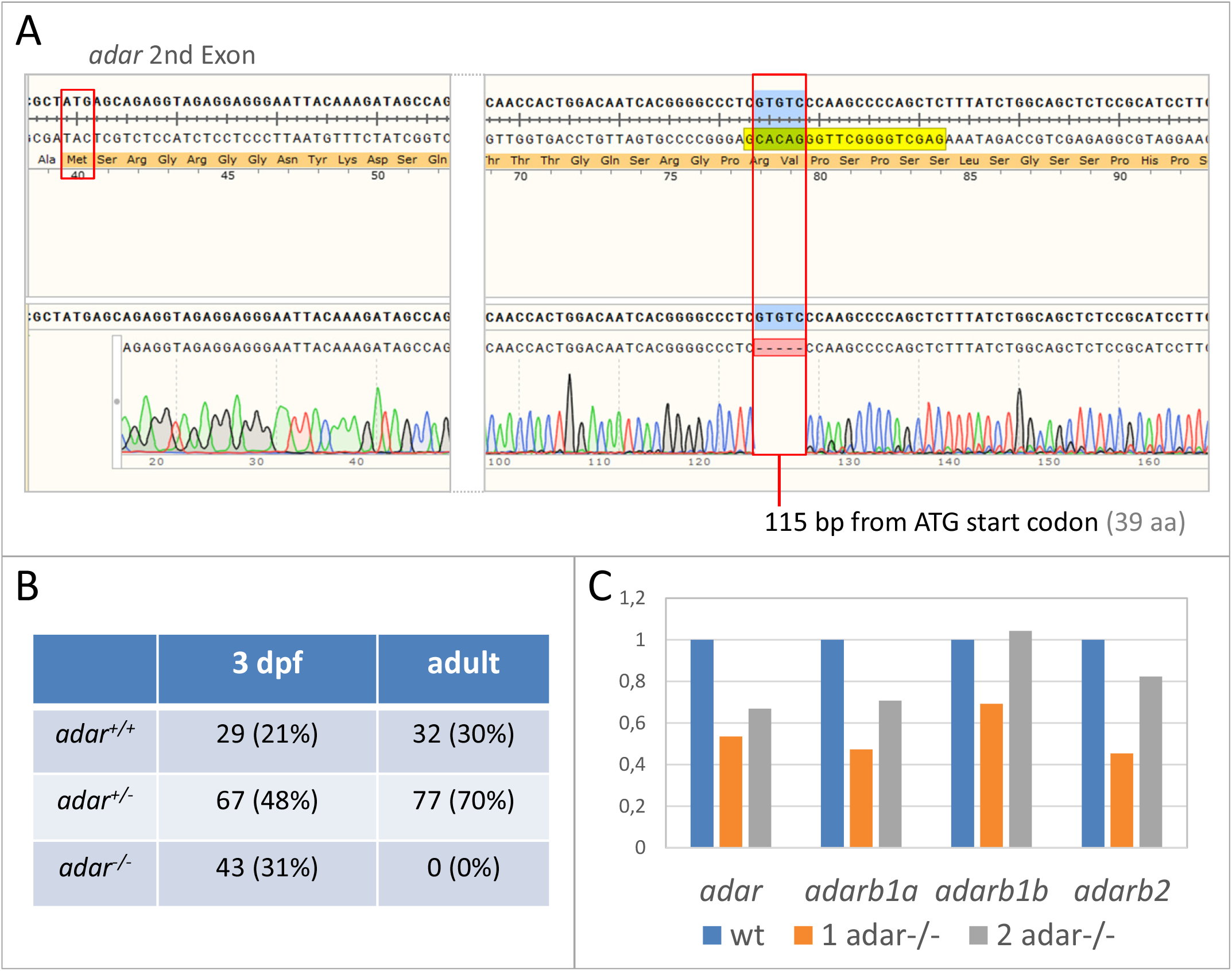
Generation of *adar-/-* mutant by CRISPR/Cas9. (A) *adar-/-* mutant had a 5-bp deletion within the second exon, resulting in a premature stop codon at 115 bp position. (B) Different genotype ratios at different time-points suggesting that *adar-/-* homozygotes die between 3 dpf and adult stage. (C) qRT-PCR of *adar* and its paralogs shows the lack of genetic compensation effect in homozygous mutant individuals.

## Discussion

### Editing primarily in the 3’-UTRs of coding transcripts

Sequencing data in this work indicate that a very small percentage of A-to-I editing in zebrafish takes place in the protein-coding regions, with the majority occurring in the 3’-UTRs of coding transcripts. Editing primarily in the 3’-UTRs and outside the coding regions of transcripts is expected based on similar patterns observed in mammals ^[51]^. It is also consistent with a recent data for zebrafish ^[23]^ that editing within the proteincoding regions of transcripts sets in after 24 hours. However, we also note that the previous study applied very conservative criteria for editing (to keep the false positive rate low) and called editing sites in embryonic transcripts based on a catalogue of editing sites for adult brain samples ^[23]^. Therefore, this study may systematically miss editing sites that are unique for the embryo and not found in the brain. Prior studies have shown that in mammals ADAR2 is more associated with editing in coding sequences, whereas ADAR1 edits more promiscuously, either in transcripts from repetitive genomic elements or, in case of coding transcripts, in the 3’-UTR regions ^[27]^. The data in this work confirm that the latter is conserved in zebrafish. Secondary structure elements located in 3’-UTR in zebrafish maternal mRNAs have been reported to be involved in translational regulation of mRNAs of the Nodal family by RNA binding proteins ^[52]^. Therefore, it is reasonable to expect that RNA editing may participate in such regulation or that similar but distinct secondary structure elements are preferred by Adar in vivo, providing a combinatorial regulatory code for fine-tuning translation and/or degradation rates. The knowledge of secondary structures adopted by maternal mRNAs in vivo is therefore becoming highly necessary to better understand the mechanism of gene expression regulation.

### No effect of Adar knockdown on transcript editing prior to the MZT

The delayed onset of embryonic transcription and reliance on maternal transcript deprives early embryos of the opportunity to regulate gene expression transcriptionally ^[35]^. This limitation, together with the very high *adar* transcript levels in early zebrafish embryos suggested to us that Adar activity could be used to control transcript and thus protein levels prior to the MZT, as an alternative to transcriptional control. Surprisingly, however, Adar knockdown or overexpression had very limited effect on global A-to-I editing until 5.3 hpf, i.e. until well after the MZT at 3.5 hpf ^[53]^. Phenotypic data and the editing data for the 12 hpf time point rule out a technical problem with knockout. It is possible (but hard to confirm in the absence of good antibodies) that a sufficient pool of maternal Adar protein is deposited in the oocyte and dominates editing in the early stages (up to 5.3 hpf), but eventually gets exhausted to reveal editing differences in the knockdown (at 12 hpf). Alternatively, editing changes as a consequence of Adar knockdown could be more subtle than anticipated. For instance, editing pattern alterations could occur within a particular transcript molecule, or different proportions of transcripts for particular genes could be edited as a result of Adar disruption. Unfortunately, the short-read sequencing method used in our study is not able to detect these differences. Long read sequencing methods such as the Oxford Nanopore promise an opportunity to further explore this in the near future.

### Different pools of Adar may explain the discrepancy between knockdown and knockout phenotypes

While the Adar KD by morpholino induced severe patterning defects, *adar-/-* mutant individuals are viable without obvious embryonic defect. Successful rescue of the knockdown phenotype with a morpholino-resistant transcript makes it unlikely that off-target effects account for the difference in editing. Genetic compensation was also ruled out in mutants. Thus, the difference in phenotype between Adar KD and mutant could be attributed to distinction of maternal and zygotic function of Adar. Instead, we suspect that maternally deposited Adar transcripts account for the difference. Due to the eventual lethality of the *adar* knockout, the *adar-/-* embryos were from *adar(+/-)* heterozygous parents. Therefore, there is a pool of maternally derived *adar* transcripts in the *adar(-/-)* embryos. By contrast, translation of the maternally derived *adar* transcripts is blocked in the morpholino experiments. We suspect that a pool of Adar proteins from embryonic translation of maternal Adar transcript spares the knockout embryo the antero-posterior and dorso-ventral defects that we see in the knockdown embryos. Eventually, however, this Adar pool is probably turned over, so that defects, now purely reflecting the embryonic genotype, become apparent. The detailed causes for this late phenotype are not yet clear. Adar null fish live vastly longer (until the second or third week) than would be expected based on death of the *adar1* null mice from hematopoietic defects (at embryonic day 12 in the mouse, equivalent to 36 hpf in zebrafish ^[54]^). Therefore, similar causes of death are unlikely. Instead, the late phenotype points to other, still unexplored roles of A-to-I editing in late larval development.

### Adar mediates embryonic patterning

Strikingly, knockdown and overexpression of *adar* resulted in opposite phenotypes, with the former abolishing posterior-ventral structures including the differentiation of notochord, while the latter affecting anterior-dorsal ones resulting in cyclopia. We also found that this phenotype is dependent on an intact RNA editing domain of Adar. These phenotypes are reminiscent of those caused by disruptions to several dorsoventral and anteroposterior axis determinants. Two signaling pathways are known to be responsible for this process: Wnt and FGF. Loss of Wnt signaling is known to cause a severe antero-dorsalized phenotype where embryos possess large heads and truncated tails, while its overactivation results in the opposite phenotype of posteriorized embryos lacking eyes ^[48,55,56]^. Similarly, loss of FGF signaling also causes truncation of the posterior body due to the lack of posterior mesoderm structures ^[57–59]^. Intriguingly, we observed that several Wnt and FGF pathway components were edited throughout the maternal and zygotic stages, suggesting a link between Adar function and the observed editing events. To our knowledge, a role of A-to-I editing in embryonic patterning is novel and has not been seen before in any vertebrate. Clearly, the very different early embryology of zebrafish compared to mammals creates opportunities for new biological outcomes of A-to-I editing that deserve further study.

## Methods

### Zebrafish

Zebrafish of wild type (AB strain) and CRISPR/Cas9-generated *adar* mutant lines were maintained in the zebrafish core facility of the International Institute of Molecular and Cell Biology in Warsaw (IIMCB) (License no. PL14656251) according to standard procedures. Embryos were raised in egg water at 28°C and staged according to standard morphological criteria^[60]^.

### MO injection, rescue and *adar* overexpression

Adar and Adarb1b knockdown was performed using a translation-blocking antisense morpholino oligonucleotides (MO) with the sequences 5′-TCCCTCCTCTACCTCTGCTCATAGC-3′ and 5’-TCCATGATGGTCAAACGTCTCGACT-3’ (Gene Tools, USA). For each embryo, 1-3 ng of MO was injected at the 1-cell stage. For overexpression experiments, the *adar* cDNA sequence was PCR-amplified using the primer pair 5’-CCTGTCTTTGATACTGTCGTG-3’ and 5’-TCCCGAAGCCACAGATTCAC-3’ and cloned into p-GEMT vector (Promega, USA). For the rescue experiment, wild-type and E1030A mutant *adar* cDNAs containing a 5 bp mismatch at the MO recognition site were PCR-amplified using the forward primer 5’-CCTAGCTAATACGACTCACTATAGGCGGAACATGAGTAGAGGAAGAGGAGGGAATTAC-3’ with a T7 promoter sequence overhang for in vitro transcription and reverse primer 5’-TCAAGCTATGCATCCAACGCG-3’. Capped mRNAs for rescue and overexpression were synthesized using the mMessage mMachine T7 Kit (Ambion, USA). Overexpression was done using 25, 50, or 100 pg of mRNA. Rescue experiments were performed using 1 ng of MO and 25 pg of wild-type or E1030A *adar* mRNA. Results were obtained from four different experiments on embryos from random pairs.

### Disruption of the deaminase domain in *adar* mRNA

Zebrafish Adar glutamate 1030 (E1030) in a highly conserved region of the protein (TVNDCHA**E**IISRRGFIRFLYSELM) was identified as equivalent to human ADAR1 glutamate 912 (E912), a residue known to be indispensable for catalytic activity of the deaminase domain. To encode the E1030A point mutation in the zebrafish *adar* clone, we used the Q5® Site-Directed Mutagenesis Kit (NEB) for mutagenic primer-directed replication of both plasmid strands. The first step was an exponential amplification using primers and a master mix formulation of Q5 Hot Start High-Fidelity DNA Polymerase. Oligonucleotides used for glutamate substitution with alanine and site-directed mutagenesis are as follows: zb_adar_E1030A_F 5’-GCAGCTATCATCTCCAGAAGAGGC−3’ and zb_adar_E1030A_R 5’-ATAGCTGCATGGCAGTCATTTACAG −3’. The second step involved an incubation with an enzyme mix containing a kinase, a ligase and *Dpn*I allowing for rapid circularization of the PCR product and removal of the template DNA. The last step was a high-efficiency transformation into chemically competent TOP10 cells. Selected clones were sent for sequencing and positive ones were linearized and used for *in vitro adar* E1030A mRNA synthesis.

### qPCR for genetic compensation analysis

Total RNA was extracted from identified (Sanger sequencing) adar KO and wild-type 5 dpf larvae, using TRIZOL LS (Thermo Fisher Scientific, USA) according to the manufacturer’s protocol, followed by DNAse I (Life Technologies, USA) treatment. Superscript IV reverse transcriptase (Life Technologies, USA) was used to obtain cDNA. Relative mRNA expression was quantified using FastStart SYBR Green master mix on the LightCycler 96 instrument (Roche Life Science, USA) with specific primer pairs (Supplementary Table 9).

### Sequencing and data analysis

For editing discovery, parental DNA was extracted from one individual male and female from tail-fin clip and sequencing library were synthesized with Nextera XT Kit (Illumina, USA) according to the manufacturer’s protocol. RNA from their offspring was also isolated at 16-cell, 3.5 hpf, and 5.3 hpf stages. For each time-point, at least 20 embryos were pooled for RNA extraction. To assess the effect of Adar KD and OE, uninjected wild-type and embryos injected with 1 ng of adar MO or 50 pg of adar mRNA were kept until the desired developmental stage: 128-cell, 50% epiboly, and 12 hpf. Two replicates of 20 pooled embryos from the first two time points and three replicates for the 12 hpf time point were isolated for RNA extraction. The RNA for the 12 hpf sample was collected in a second round of experiments, from offspring of a separate mating pair. Total RNA was extracted using TRIzol LS (Thermo Fisher Scientific, USA) and cleaned up on the Qiagen Rneasy Mini column (Qiagen, USA). Quality control of extracted RNA was performed using the 2200 TapeStation system from Agilent Technologies (USA). To avoid the bias caused by cytoplasmic polyadenylation during early embryonic stages, polyA affinity was not used for mRNA enrichment. Instead, total RNA was rRNA-depleted using Ribo-Zero Magnetic Gold Kit (Human, Mouse, Rat; Epicenter). cDNA synthesis for Next-Generation Sequencing (NGS) was performed by SMARTer Stranded RNA-seq kit (Clontech Laboratories, USA) as recommended by the manufacturer. Paired-end sequencing (2 × 75 bp reads) was performed with NextSeq 500 (Illumina, USA). Reads were aligned to the zebrafish genome assembly GRCz10 using STAR v2.7.7a ^[61]^ and samtools v1.11 ^[62]^. On average, more than 80% of total sequencing reads were uniquely mapped (Supplementary Table 10). Expression quantification was performed using HTSeq v0.11.2 ^[63]^. Differential expression was performed using DESeq2 (R v3.6.3) ^[64]^. HTseq reads from KD and OP samples were compared to the control samples at corresponding time points. Multiple testing was done by applying the Benjamini-Hochberg correction as implemented in DESeq2 with adjusted p-values <0.05 called as statistically significant.

### RNA Editing Discovery

Putative RNA editing sites based on DNA- and RNA-seq input were detected using REDiscover (https://github.com/lpryszcz/REDiscover). The script automatically eliminates low quality and duplicate reads. It utilizes samtools mpileup to generate a text file from input bam files. Options-q 15-Q 20, were used, i.e. a minimal mapping quality of 15 and a minimal base call quality of 20 were required. Sites that were not homozygous between female and male samples were excluded from the analysis. Alternative alleles were only called when they were present in at least 20% of RNA reads. For simplicity, sites with more than one alternative allele were excluded from the analysis. Between 57,605 and 150,495 putative edited sites were detected per sample (Supplementary Table 1). These sites were then filtered in a post-processing step that required a minimum coverage of 10 reads in both the DNA-Seq and RNA-Seq data.

### Whole-mount *in situ* hybridization

Whole-mount *in situ* hybridization was performed on 24 hpf wild-type, *adar* morpholino-injected and *adar* mRNA-injected embryos. Dechorionated embryos were fixed in 4% PFA for 2 hours at room temperature, washed in PBS/Tween (PBT) and then sequentially dehydrated in methanol. After 100% methanol overnight incubation at − 20°C, the embryos were rehydrated in serial methanol dilutions, washed in PBT, digested with 10 μg/mL Proteinase K (Roche) for 5 min and refixed in 4% PFA for 20 min at room temperature. Then, they were washed in PBT and incubated in hybridization buffer (Hyb; 5x SSC, 50 μg/ml heparin, 0.1% Tween 20, 500 μg/mL tRNA, 50% formamide) in a water bath at 68°C. After 4 hours, the embryos were incubated in hybridization buffer containing 1 μg/mL DIG-labelled RNA probe overnight at 68°C. Thereafter we performed several post hybridization washes: serial Hyb/2× SSC dilutions for 10 min each at 68°C, 2×10 min 2× SSC at 68°C, 2x15 min 0.2× SSC at 70°C and serial 0.2× SSC/PBT dilutions at room temperature. The embryos were incubated in blocking solution (2 mg/mL BSA, 2% sheep serum and 1× PBT) at room temperature for 3-4 hours, after which they were incubated in anti-DIG antibody (Roche, 1:5000) in the dark at 4°C overnight. They were washed several times in PBT and incubated in alkaline phosphatase buffer (NTMT; 0.1 M Tris-HCl pH 9.5, 50 mM MgCl_2_, 0.1 M NaCl, 0.1% Tween 20) 3 times for 5 min each. We then stained the embryos by incubating them with NBT/BCIP solution (Roche) at room temperature. When the desired staining intensity was reached, the reaction was stopped by washing the embryos in PBT and fixing them in 4% PFA. Pictures were taken using Nikon SMZ25 stereomicroscope.

The *adar, adarb1a, adarb1b, gsc, foxa2, shhb* and *tbxta* clones were amplified from a cDNA template using specific primers containing a T7 promoter sequence overhang at the reverse primer (Supplementary Table 9), and the corresponding riboprobe was synthesized using the DIG-RNA labeling kit (Roche) according to the manufacturer’s instructions. Clones for *pax6* and *tbx2b* were a kind gift from Vladimir Korzh.

### CRISPR/Cas9-mediated *adar* knock-out in zebrafish

sgRNAs targeting the *adar* gene were designed using the CCTop on-line tool ^[65]^. Three potential sequences were chosen based on their vicinity to the start codon as well as lack of predicted off-target exonic sites. Each sgRNA was tested by co-injecting it with Cas9 mRNA into 1-cell stage embryos. At 24hpf genomic DNA was extracted from single embryos using the HOTSHOT method ^[66]^. Genomic DNA was then used as a template for HRM analysis using sets of primers specific to the targeted region of the gene. DNA from single uninjected embryos was used as negative control. Based on the highest percentage of edited embryos, sgRNA_3 (5’-GAGCTGGGGCTTGGGACACG - 3’) was deemed the most efficient and used for subsequent knock-out line generation. In order to establish the mutant line, embryos of wild-type (AB) line fish at 1-cell stage were injected with 40 pg sgRNA_3 and 400 pg Cas9 mRNA each. The embryos were then raised to adulthood and outcrossed with wild-type AB fish. DNA from embryos resulting from this spawning was extracted and analyzed using the HRM approach as described before to screen for germline transmission ^[67]^. Outcross of edit-carrying F0 individuals with wild type resulted in 25% of heterozygotes (F1). Offspring of incrossed F1 heterozygotes were raised to three months post fertilization and subsequently genotyped to identify mutant homozygotes (*adar* -/-). Oligonucleotides used for genotyping are as follows: adar_genom_F 5’-CTAACGCTACACCCTCCTCAGC-3’ and adar_genom_R 5’-CGTCTGGTACTGGATAGGCTC −3’. Similarly, 3 dpf larvae from F2 generation were genotyped according to a published protocol ^[68]^ and sequenced using the abovementioned pair of oligonucleotides.

## Data availability

All sequencing data have been deposited in the GEO database under accession number GSE182714.

## Authors’ contributions

C.W. and M.B. conceived the project and designed the experiments. K. N., C.N.H., E.T., C.W., and M.K. performed experiments. K.A.N. and K.M. prepared libraries and performed sequencing. L.P. developed analyses pipelines and performed data analysis with contribution from M.B. C.W., M.B., K.N., and E.T. wrote the manuscript. All authors have read and approved the final manuscript.

## Acknowledgements

We are grateful to the zebrafish core facility of the IIMCB Warsaw for excellent fish care; S. Guenther and T. Braun for permission to use sequencing facility and assistance in sequencing runs; M. Łapiński and M. Migdał for assistance in data transfer. We thank M. O’Connell, L. Solnica-Krezel, V. Korzh, as well as members of the C.W. and M.B. labs for fruitful discussions.

## Suppl. Figure Legends

**Supplementary Figure 1.**
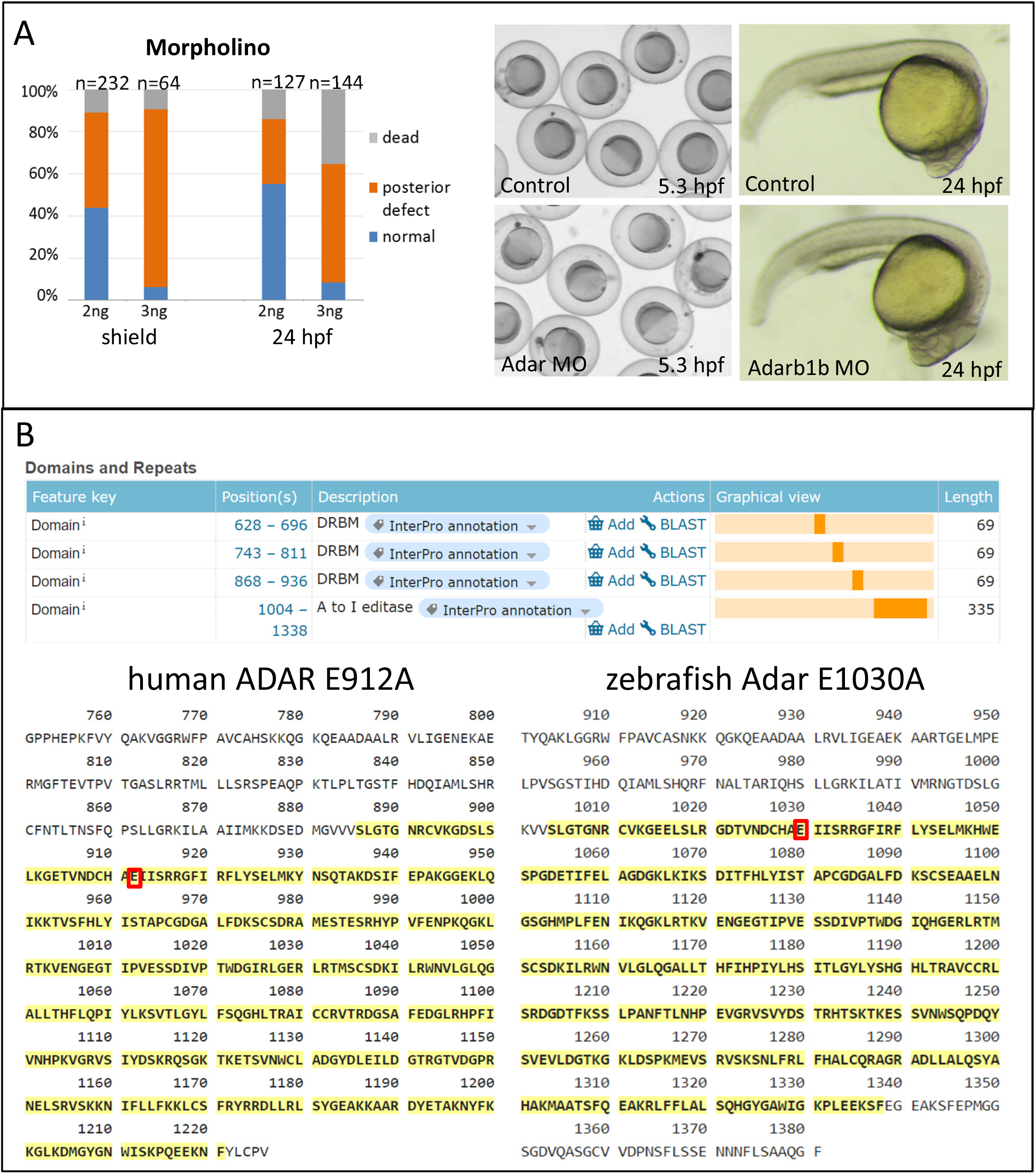
Adar and Adarb1b knockdown experiments. **(A)** Dose-dependent effect of Adar MO was observed starting from 5.3 hpf, while Adarb1b MO did not cause any observable phenotype. (B) Design of *adar* mutant mRNA E1030A with point mutation abolishing the activity of the deaminase domain.

**Supplementary Figure 2.**
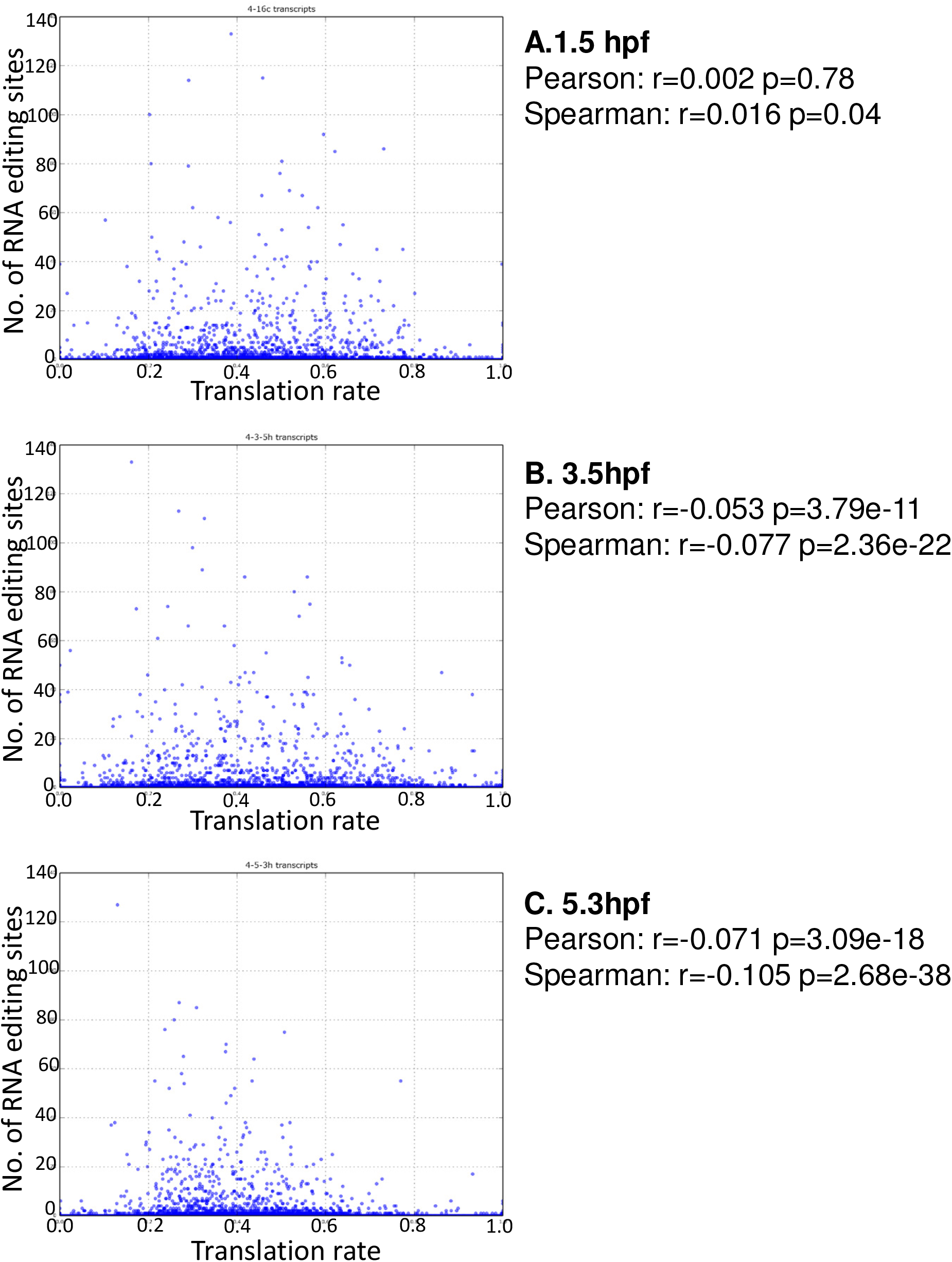
Translation rates and number of editing sites for 1.5 hpf and 5.3 hpf transcripts. Translation rates are expressed as ratio of polysome bound to sum of bound and unbound fractions of a given transcript ^[54]^. Thus, translation rate of 1.0 means all expressed transcript molecules are associated with polysome, while 0.0 means none of expressed transcript molecules are associated with polysome.

**Supplementary Figure 3.**
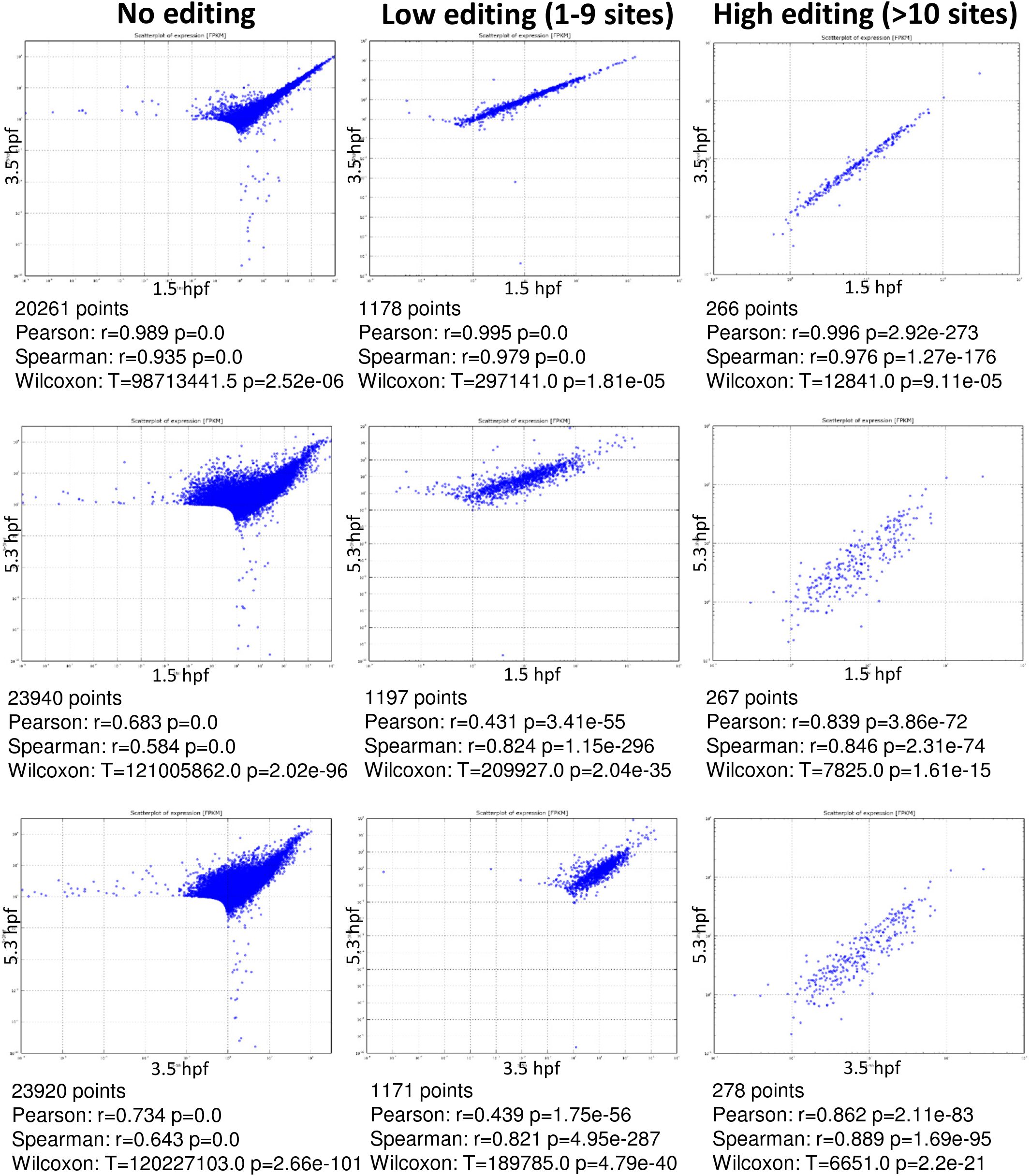
Comparison of expression levels between two developmental stages of non-edited, low-edited, and highly edited transcripts.

**Supplementary Figure 4.**
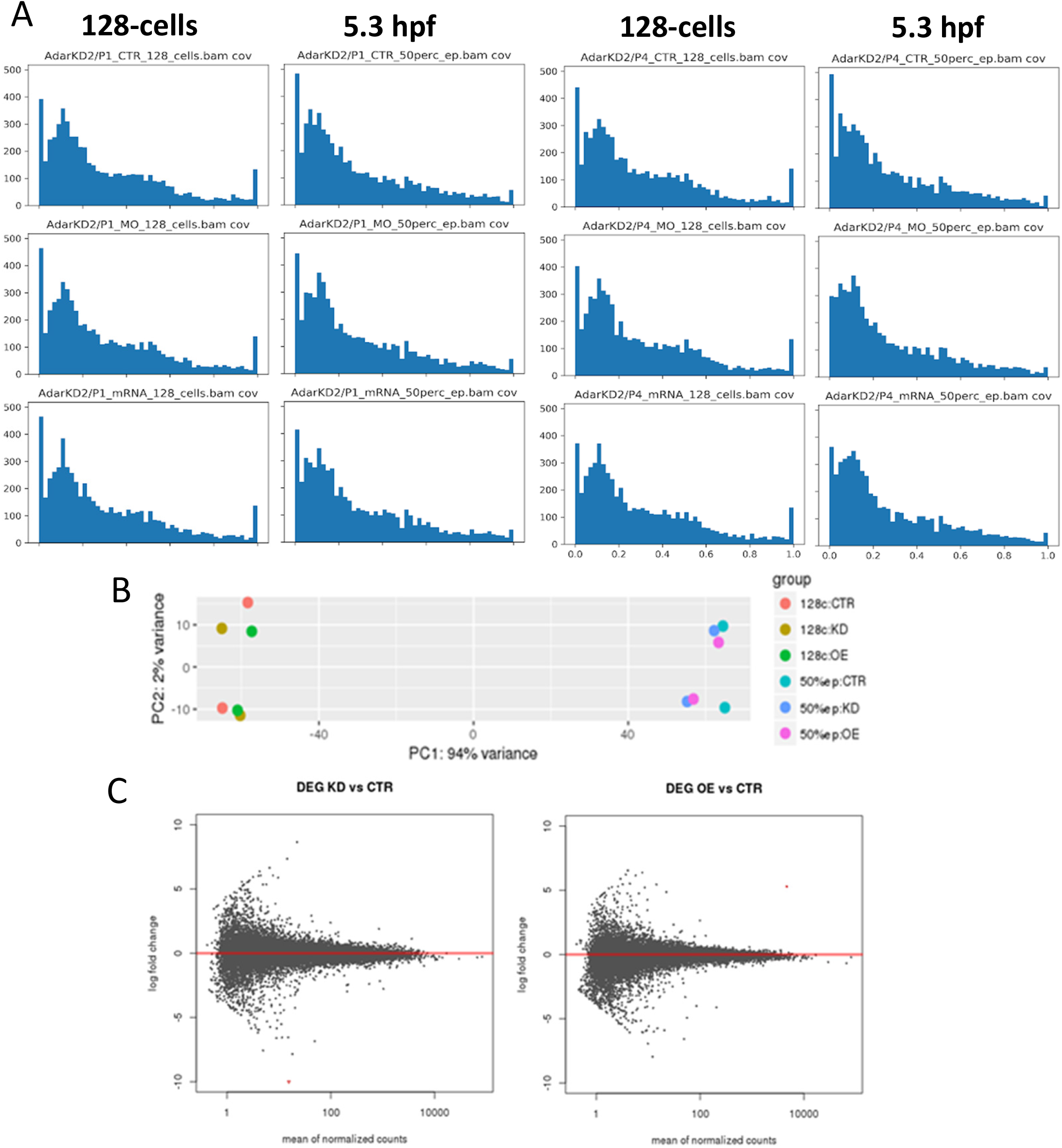
Global RNA editing and gene expression profile of Adar knockdown and overexpression. (A) Penetrance of RNA editing expressed as number of transcripts vs. fraction of editing within a transcript species in 128-cell (left column) and 5.3 hpf (right column). (B) Principal component analysis of control, Adar knockdown and overexpression samples based on their transcriptome profile. (C) Differential expression analysis of Adar knockdown and overexpression vs. control. No genes were differentially expressed at p < 0.05.

**Supplementary Figure 5.**
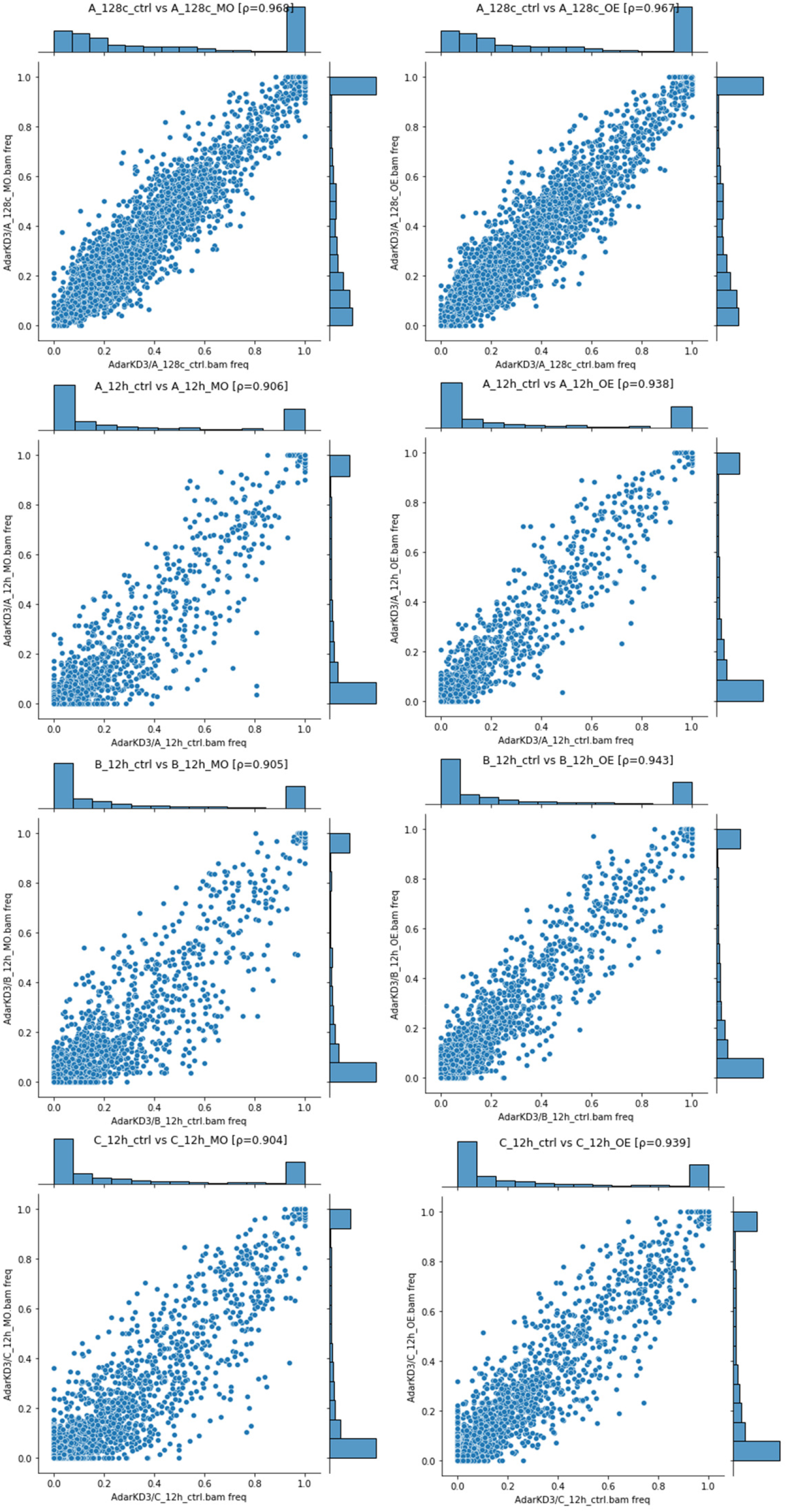
Comparison of RNA editing frequency between control and Adar KD or OE. RNA editing frequencies for each transcript is plotted for each sample in 128-cell and 12 hpf stages. Spearman’s rank correlation coefficient (ρ) is given in every figure title.

